# Aging enteric neurons develop chromatin remodeling, DNA damage and a senescence-like phenotype

**DOI:** 10.64898/2026.11.11.765843

**Authors:** Lobke Mombeek, Anton Brosens, Reindert Jehoul, Anita J. L. Leembruggen, Peter Neckel, Marlene Hao, Werend Boesmans, David M Wilson

## Abstract

Aging is associated with gastrointestinal dysfunction, yet the cellular and molecular changes underlying aging of the enteric nervous system (ENS), the cellular network that controls gut motility and digestion, remain poorly characterized. Here, we investigated whether enteric neurons show canonical hallmarks of aging in mice and humans. We assessed age-related changes in enteric neurons of the jejunum and colon in mice aged 2, 13-15, and 22-24 months using immunofluorescence, transmission electron microscopy, Ca²⁺ imaging, and network morphology analysis, and compared selected markers between young and aged human gut tissue. Enteric neurons accumulated DNA damage with age in both the jejunum and colon, with nitrergic neurons exhibiting a greater burden of damage than cholinergic neurons. Increased H3K9me3-DAPI colocalization, consistent with senescence-associated heterochromatin foci, was already seen at 15 months in both regions, revealing a potentially early driver of age-related gut changes. At 24 months, colonic neuron numbers decreased, while average neuronal soma size was increased and approximately 25-30% of neurons expressed cytoplasmic p21, a marker of senescence. Aged neurons showed enlarged, less circular mitochondria and increased pPDH immunoreactivity, indicative of declining neuronal activity. Consistent with remodeling of myenteric plexus architecture, neuronal Ca²⁺ responses increased upon electrical stimulation while decreasing after agonist application. DSS-induced colitis did not reproduce the DNA damage, heterochromatin or p21 changes seen with age, implying a direct role in the latter. Human aged enteric neurons showed higher γH2AX and H3K9me3 intensity. These data demonstrate that enteric neurons develop a senescence-like phenotype with age, and identify neuronal senescence as an underexplored aspect of ENS aging.

## INTRODUCTION

The enteric nervous system (ENS) is a gut-intrinsic network of neurons and glia that autonomously regulates major aspects of gastrointestinal (GI) function (1), such as bowel motility, mucosal secretion, and blood flow (2,3). Owing to its size and close anatomical proximity to the gut lumen, the ENS is uniquely positioned among neural systems to encounter both endogenous and exogenous stressors across the lifespan. Enteric neurons and glia are continuously exposed to natural intracellular reactive chemical species, as well as to fluctuations in local inflammatory cues and metabolic stress generated by ongoing digestive and immune activity (4–6). Additionally, the ENS continuously encounters potential external environment threats via access through the gut wall, such as including dietary components, ingested toxins, medications, and metabolites from shifts in the gut microbiota and its metabolites, throughout an individual’s lifetime (7,8). This sustained, cumulative exposure suggests that ENS integrity may reflect not only an individual’s intrinsic physiological stress burden but also the sum of environmental influences encountered over a lifetime, making it a particularly compelling system for studying the biological consequences of aging.

Aging is characterized by the gradual decline in tissue and organ functionality and is associated with several well-established hallmarks (9). Primary hallmarks, which encompass core molecular and cellular alterations that likely drive aging, include genomic instability, telomere attrition, epigenetic alterations, and loss of proteostasis. These triggers are presumed to underlie downstream antagonistic hallmarks, namely mitochondrial dysfunction and cellular senescence, and integrative hallmarks, such as stem cell exhaustion and altered intercellular communication. Different organs manifest aging hallmarks in very distinct ways, implying distinct aging profiles that depend on cell- and organ-type and their personalized “experiences” (10,11). While much analysis has gone into characterizing these hallmarks in a wide-range of cell-types, tissues and organs, very little is known with respect to the ENS.

Early studies have reported that morphological changes in the aging ENS likely affect gut motility, secretion, and sensation. Specifically, the general neuronal loss and reorganization of myenteric ganglia seen in guinea pigs would disrupt neural control, impairing GI function (12). In rats, dystrophic features, such as swollen axons and terminals, point to structural damage that would likely compromise neuronal signaling (13). Being vital for gut function, the emergence of aging hallmarks within the ENS would most certainly contribute to the development of GI dysfunction, culminating in problems like dysphagia, gastroesophageal reflux, constipation, and fecal incontinence, symptoms known to become more prevalent with age (14–17).

To date, Jurk et al. have shown that mouse enteric neurons accumulate DNA damage with aging and exhibit signs of senescence, such as increased senescence-associated β-galactosidase activity, activated p38 mitogen-activated protein kinase (MAPK), and elevated levels of oxidative stress (18). More recently, Palmer et al. extended this observation to human tissue, reporting increased expression of the senescence marker p16 within myenteric neuron cell bodies of the aged human ascending colon, suggesting that a comparable post-mitotic senescence-like phenotype may occur (19). Enteric glia, which support neuronal survival and function, also decline in number with advancing age (20), indicating that aging affects the ENS beyond its neuronal population alone. More broadly, age-related ENS dysfunction may be compounded by changes in the gut’s resident immune environment: self-maintaining populations of intestinal macrophages positioned close to myenteric and submucosal ganglia are required for enteric neuron maintenance, and their depletion leads to enteric neuron loss and impaired motility (21). The collective molecular and clinical studies suggest that enteric neurons are vulnerable to age-associated genomic and cellular stress, and that this susceptibility, potentially exacerbated by a decline in glial and immune support, may contribute to GI dysfunction observed with aging.

To more thoroughly define the molecular drivers underlying ENS aging, we have herein characterized the aging phenotype of enteric neurons across mouse and human tissue by combining immunofluorescence labeling of markers of key aging hallmarks, i.e., genomic stress, epigenetic alterations, mitochondrial integrity, metabolic activity, and senescence, with Ca²⁺ imaging and electron microscopy. We show that aging enteric neurons in both species accumulate DNA damage and adopt a more pronounced senescent phenotype than previously recognized. Importantly, chromatin remodeling emerges at midlife, preceding the broader manifestation of aging-associated neuronal alterations, suggesting that epigenetic dysregulation may represent an early event and potential driver of ENS aging. Together, these findings position enteric neuron senescence and epigenetic remodeling as underappreciated, conserved features of ENS aging across species.

## MATERIALS AND METHODS

### Animals

A total of 48 C57BL/6J mice were used in this study, with males and females equally represented across three age groups: 2 months, 13-15 months, and 22-24 months. Animals were housed under standard conditions on a 12 h:12 h light–dark cycle in temperature-controlled rooms (20 °C ± 3 °C) with ad libitum access to autoclaved food and water. All mice were maintained in filter-top cages and provided with appropriate environmental enrichment. Mice were sacrificed by cervical dislocation. All animal procedures were performed in accordance with Belgian regulations and were approved by the Ethical Committee for Animal Welfare of Hasselt University.

### DSS Treatment

A separate cohort of four male mice was used for the dextran sulfate sodium (DSS) challenge described in Figure 5. Animals were 3 months of age at the start of treatment. Acute colitis was induced by administering 3% DSS (colitis grade, 36-50 kDa; MP Biomedicals, 02160110-CF) in drinking water ad libitum for 7 consecutive days prior to sacrifice. Animals were monitored daily for body weight, stool consistency, rectal bleeding, general appearance, and overall welfare, and these parameters were used to assess disease progression using the Disease Activity Index (DAI). At the end of the treatment period, mice were sacrificed by cervical dislocation. To confirm the induction of acute colitis, hematoxylin and eosin (H&E) staining was performed on 4-10 μm sections of paraffin-embedded small and large intestinal tissue fixed in 4% paraformaldehyde (PFA). Whole-slide images were acquired using a Pannoramic 480 Digital Scanner (3DHISTECH). All procedures were performed in accordance with Belgian regulations and were approved by the Ethical Committee for Animal Welfare of Hasselt University.

### Immunofluorescence in mouse PFA-fixed tissues

Full-length gut was immersed in ice-cold 1× PBS (137 mM NaCl, 2.7 mM KCl, 10 mM Na₂HPO₄, 1.8 mM KH₂PO₄), opened along the mesenteric border, and pinned with the epithelium side facing up on plates coated with Sylgard 184 Elastomer (Dow, catalog number 01673921) using insect pins. The tissue was then fixed with 4% PFA in 1× PBS for 2 hours at room temperature. Following fixation, the mucosa and circular muscle layers were peeled off, and the resulting longitudinal myenteric plexus muscle preparations were cut into 1 cm² sections and stored in 0.01% sodium azide in 1× PBS at 4°C for both colonic and jejunal tissue. Prior to staining, the stored tissues were rinsed three times for 5 minutes each with 1× PBS. For antigen retrieval, a solution was prepared by diluting HistoVT One buffer (Nacalai Tesque, catalog number 06380-05) 1:10 in Milli-Q water. This antigen retrieval step was performed at 70°C for 20 minutes for γH2AX or p21 stainings. Permeabilization was achieved by adding 0.1% Tween-20 to 1× PBS, and the blocking solution consisted of 10% goat serum (Millipore, catalog number 4064259) or donkey serum (Millipore, catalog number S30-100ML) and 0.1% Triton X-100 in 1× PBS. Tissue samples were washed three times for 10 minutes each with 1× PBS. Primary antibody incubation was performed by incubating the tissues with the primary antibody solution in blocking buffer overnight at 4°C (Table 1). After primary antibody incubation, tissues were rinsed three times for 10 minutes each with 1× PBS. Secondary antibody incubation was carried out by incubating the tissues with the secondary antibody solution in blocking buffer for 2 hours at 4°C (Table 1). Following secondary antibody incubation, tissues were washed three times for 10 minutes each with 1× PBS and then incubated with 1:10000 4′,6-diamidino-2-phenylindole (Thermo Fisher Scientific, catalog number 62248) in 1× PBS. Finally, the tissues were washed three more times for 10 minutes each with 1× PBS and mounted using Fluoromount-G mounting medium (Thermo Fisher Scientific, catalog number 00-4958-02).

**Table 1.** Primary and secondary antibodies used.

| **Primary antibodies** | **Dilution** | **Manufacturer** | **RRID** |
|---|---|---|---|
| Human anti-HuC/D | 1:20000 | Gift from Dr. Vanda Lennon, Mayo Clinic |  |
| Mouse anti-HuC/D | 1:200 | Thermo Fisher Scientific, A-21271 | AB_221448 |
| Rabbit anti-p21 | 1:500 | ProteinTech, 10355-1-AP | AB_2077682 |
| Rabbit anti-γH2AX | 1:800 | Cell Signalling, 9718 | AB_2118009 |
| Sheep anti-nNOS | 1:3000 | Gift from Dr. Piers Emson, Cambridge University |  |
| Goat anti-ChAT | 1:100 | Millipore, AB144P | AB_2079751 |
| Rabbit anti-H3K9me3 | 1:1000 | Cell Signalling, 13969 | AB_2798355 |
| Rabbit anti-pPDH | 1:1000 | Cell Signalling, 37115 | AB_2923272 |
| Rabbit anti-PGP9.5 | 1:400 | Proteintech, 14730-1-AP | AB_2210497 |
| **Secondary antibodies** | **Dilution** | **Manufacturer** |  |
| Goat anti-Human 555 | 1:1000 | Invitrogen, A-21433 |  |
| Goat anti-Rabbit 555 | 1:1000 | Invitrogen, A-21430 |  |
| Goat anti-Rabbit 647 | 1:1000 | Invitrogen, A-21245 |  |
| Donkey anti-Goat 488 | 1:1000 | Invitrogen, A-11055 |  |
| Donkey anti-Sheep 488 | 1:1000 | Invitrogen, A-11015 |  |
| Donkey anti-Rabbit 647 | 1:1000 | Invitrogen, A-31573 |  |
| Goat anti-Mouse 488 | 1:1000 | Invitrogen, A-21141 |  |
| Goat anti-Rat 647 | 1:1000 | Invitrogen, A-21247 |  |

### Immunofluorescence in human paraffin-embedded gut sections

Human duodenal samples were collected from 4 XX female and 2 XY male patients aged between 14 and 82 (Table 2). All samples were obtained at the University Hospital Tübingen, Germany after approval by the local ethical committee (Project Nr. 652/2019BO2 and 066/2023BO2), and according to the declaration of Helsinki. Paraffin-embedded tissue sections were deparaffinized in xylene and graded ethanol solutions before rehydration in distilled water. Heat-induced antigen retrieval was performed in citrate buffer (10× stock: 29.5 g/L trisodium citrate dihydrate, 0.5% Tween-20, pH 6.0) using microwave heating. Sections were subsequently washed in PBS and permeabilized in PBS containing gelatin and 0.5% Triton X-100. Non-specific binding was blocked with 5% goat serum for 1 h at room temperature. Primary antibodies were diluted in permeabilization buffer containing 1% goat serum and incubated overnight at 4°C (Table 1). After washing, sections were incubated with species-appropriate fluorescent secondary antibodies (1:1000) for 2 h at 4°C (Table 1). Nuclei were counterstained with DAPI (1:10,000) for 10 min at room temperature. Finally, sections were mounted using Fluoromount-G (Thermo Fisher Scientific, 00-4958-02) mounting medium and allowed to cure before imaging.

**Table 2.** Details on human duodenal samples.

| **Group** | **Age** | **Sex** |
|---|---|---|
| Young | 14 years and 7 months | Female |
| Young | 19 years | Female |
| Young | 27 years | Female |
| Age | 63 years | Male |
| Age | 67 years | Female |
| Age | 82 years | Male |

### Widefield, confocal and multiphoton microscopy

Samples for mouse HuC/D neuron counting, pPDH staining (mouse and human), and PGP9.5 staining were imaged using a Nikon Eclipse Ti2-E fluorescence microscope equipped with a Kinetix sCMOS camera and a CoolLED pE800 light source (550 LED-DA/FI/TR/Cy5/Cy7) for red and far-red fluorescence detection. Imaging was performed using a 20× CFI Plan-ApoChromat Lambda objective lens (Nikon). Mouse γH2AX and p21, as well as human γH2AX and H3K9me3 stainings, were acquired using a Zeiss LSM900 Airyscan 2 confocal microscope equipped with CW 488 nm, CW 561 nm, and CW 640 nm diode lasers and a Plan-ApoChromat 20×/0.80 M27 objective lens. H3K9me3 staining was additionally imaged using a Leica DM4000 B LED microscope equipped with a DFC450C color camera, appropriate red and green filter sets, and 20×/40× objective lenses. Second harmonic generation (SHG) imaging was performed using a Zeiss LSM 880 IndiMo multiphoton microscope (Axio Observer platform) equipped with a Ti:Sapphire laser tuned to 810 nm and non-descanned detectors. Images were acquired using an EC Plan-Neofluar 10×/0.30 air objective (Zeiss). Tile-scan images were collected at a resolution of 512 × 512 pixels per tile and subsequently stitched using the Zeiss acquisition software. Identical acquisition settings, including laser power, detector gain, scan speed, and objective configuration, were maintained across all samples to ensure comparability between experimental groups. Representative images were uniformly adjusted for brightness and contrast using ImageJ.

### Image analysis and statistics

Image analysis was performed using Fiji/ImageJ (22). Cell quantification was carried out manually using the Cell Counter plugin unless otherwise specified. For γH2AX analysis, images were separated into individual channels, after which neuronal nuclei were identified from the HuC/D channel using Gaussian blurring (radius = 3), Otsu thresholding, binarization, and watershed separation. Total neuronal nuclei per field of view were quantified using the Analyze Particles function. DNA damage was subsequently assessed in the γH2AX channel, and neurons were manually classified as γH2AX-negative (<5 foci) or γH2AX-positive (≥5 foci). The proportion of γH2AX-positive neurons was calculated relative to the total number of segmented neuronal nuclei. For p21 analysis, the number of positive cells was normalized to the total number of neurons identified using HuC/D, nNOS, or ChAT markers. For H3K9me3 analysis, myenteric ganglia were manually delineated based on the HuC/D signal, after which colocalization between the H3K9me3 and DAPI channels was quantified using the Coloc2 plugin. Pearson’s correlation coefficient (Pearson’s R) was calculated for each ganglion and used as a measure of H3K9me3-DNA colocalization. In human samples, mean fluorescence intensity (MFI) was quantified following manual ROI selection based on HuC/D staining. SHG signal was quantified by manually delineating the muscle layer using tissue autofluorescence to define regions of interest (ROIs). The SHG channel was thresholded (cut-off = 30), binarized, and the SHG-positive area within each ROI was measured. Data were compiled in Microsoft Excel and analyzed using GraphPad Prism 9.5.1. Data are presented as mean ± SD. Normality was assessed using the Shapiro-Wilk test. For normally distributed data, independent t-tests, one-way ANOVA with Tukey’s post hoc test, or two-way ANOVA were applied. Non-normally distributed data were analyzed using Mann-Whitney or Kruskal-Wallis tests with Dunn’s post hoc correction. Statistical significance was defined as p < 0.05 (p < 0.05, p < 0.01, p < 0.001, p < 0.0001). Unless otherwise stated, these statistical approaches were applied throughout the study; any deviations are specified in the corresponding figure legends.

### Quantification of myenteric plexus network morphology

Quantitative analysis of PGP9.5-immunoreactive myenteric plexus networks was performed in Fiji/ImageJ. Prior to analysis, images were subjected to background subtraction using the rolling-ball algorithm, followed by noise reduction using a median filter (radius = 5 pixels). Segmentation of the neuronal network was achieved using automated Triangle thresholding, after which binary masks were generated for subsequent analyses. Network density was quantified by measuring the area fraction (%) of the thresholded image, representing the proportion of the field of view occupied by PGP9.5-positive structures. Fiber bundle thickness was assessed using the Local Thickness plugin in Fiji. A cutoff value of 30 was applied to exclude small structures and image noise. Local thickness values were exported and represented as frequency histograms using 50 bins to evaluate the distribution of fiber bundle diameters within the network. To assess network architecture, binary images were skeletonized and analyzed using the Analyze Skeleton plugin. Branch lengths identified by the software were extracted, and all non-zero branch lengths were included in the analysis. The average branch length was subsequently calculated and used as a measure of neuronal network organization and complexity. All image processing and analysis steps were performed using identical parameters for all samples.

### Transmission electron microscopy

Whole-gut tissue samples were fixed in Karnovsky’s fixative (2.5% glutaraldehyde and 2% paraformaldehyde in 0.1 M sodium cacodylate buffer, pH 7.4). Following fixation, samples were washed five times in 0.1 M sodium cacodylate buffer and post-fixed in 1% osmium tetroxide supplemented with 1.5% potassium ferricyanide in the same buffer for 1 h at 4 °C in the dark. After postfixation, samples were rinsed five times in Milli-Q water and dehydrated through a graded ethanol series (70% ethanol, 2 × 30 min; 90% ethanol, 2 × 30 min; 100% ethanol, 3 × 30 min). This was followed by three incubation steps in 1,2-propylene oxide (3 × 30 min). For resin infiltration, tissues were first incubated overnight in a 1:1 mixture of epoxy resin and ethanol, followed by incubation in pure epoxy resin for 6 h. Samples were then embedded in fresh epoxy resin and polymerized for >48 h at 60 °C. Ultrathin sections (70 nm) were cut using a Leica Ultracut UCT ultramicrotome and mounted on Formvar-coated copper grids. Sections were stained with 2% uranyl acetate in 50% ethanol followed by lead citrate. Imaging was performed using a Tecnai T12 transmission electron microscope equipped with an Eagle 4 k × 4 k CCD camera (Thermo Fisher Scientific).

### Ca^2+^-imaging in live muscle myenteric plexus preparations

For calcium imaging experiments, adult Wnt1::Cre;R26R-RCL-GCaMP3 mice (hereafter referred to as Wnt1-GCaMP3 mice) were utilized. In this model, the genetically encoded calcium indicator GCaMP3 is selectively expressed in neural crest-derived cells, including all ENS populations. Wnt1-GCaMP3 mice were generated by crossing Wnt1::Cre mice (23) with R26R-RCL-GCaMP3 (Ai38; The Jackson Laboratory, stock no. 029043) reporter mice (24). To minimize any neural plasticity and gut motility changes due to the estrus cycle (25), only male mice were used in this study. All experimental procedures were approved by the Animal Ethics Committee of the University of Melbourne (No. 1714226 and No. 21474). Live Ca²⁺ imaging was carried out on ex vivo preparations of the myenteric plexus isolated from the proximal colon. Dissections and all recordings were performed in Krebs’ solution containing (mM): 117 NaCl, 4.6 KCl, 2.5 CaCl₂, 1.2 MgSO₄, 1 NaH₂PO₄, 25 NaHCO₃, and 11 D-glucose. Colons were pinned in Sylgard 184–coated Petri dishes (Dow Corning) and opened along the mesenteric border. The mucosal layer, submucosal plexus, and portions of circular muscle were removed to obtain a longitudinal muscle–myenteric plexus (LMMP) preparation, which was mounted on a small stainless-steel ring for imaging, as previously described (26). Preparations were imaged in glass-bottom Petri dishes (Ibidi) using an inverted microscope (Axiovert 25, Zeiss) equipped with a ×20 (NA 0.5) objective and an Axiocam 702 mono camera (Zeiss). GCaMP3-positive cells were excited with a 470 nm LED light source (Zeiss Colibri), with exposure times of 30–50 ms. Time-lapse recordings were acquired for 1 min at 2 Hz (512 × 512 pixels). Following an initial recording of baseline activity for 10 sec, neuronal activity was evoked with focal electrode stimulation (Tungsten wire 50µm diameter; 20 pulses, 300µsec, 20Hz) 1,1-dimethyl-4-phenylpiperazinium (DMPP, 10µM), or ATP (10µM) (27). Fluorescence changes of GCaMP3, reflecting intracellular Ca²⁺ dynamics, were recorded using Zen Pro software (Zeiss). Data analysis was performed in Igor Pro (WaveMetrics) using custom-written scripts (available via https://gbiomed.kuleuven.be/english/research/50000625/50000628/site/lens.htm) (28). Regions of interest were manually selected, and fluorescence intensity was quantified per cell and normalized to baseline fluorescence (mean of the first 10 frames), expressed as F/F₀. Each pharmacological condition was recorded in preparations obtained from at least three mice.

## RESULTS

### Enteric neurons accumulate DNA damage and exhibit epigenetic alterations with age

To assess DNA damage, a driver of the primary “genomic instability” hallmark, γH2AX immunostaining was performed on the myenteric plexus of young (2 mo), adult (15 mo), and aged (24 mo) mouse jejunum and colon. Because γH2AX forms discrete nuclear foci upon phosphorylation at sites of DNA double-strand breaks, we established a scoring method in which neurons containing more than five γH2AX foci were classified as γH2AX-positive (Fig. 1A, Supplementary fig. 1A). γH2AX and HuC/D (pan-neuronal marker) co-staining in young and aged colonic LMMPs (Fig. 1B) showed a subtle but consistent increase in γH2AX accumulation with age. Quantification confirmed a significant age-dependent rise in γH2AX-positive neurons in both the jejunum and colon (Fig. 1C). To determine whether specific neuronal subtypes are differentially susceptible to DNA damage, we co-labelled γH2AX with ChAT or nNOS (Fig. 1D). In the jejunum, nitrergic neurons exhibited significantly higher γH2AX levels than cholinergic neurons, whereas no significant difference between the two subtypes was observed in the colon (Fig. 1E).

**Figure 1.**
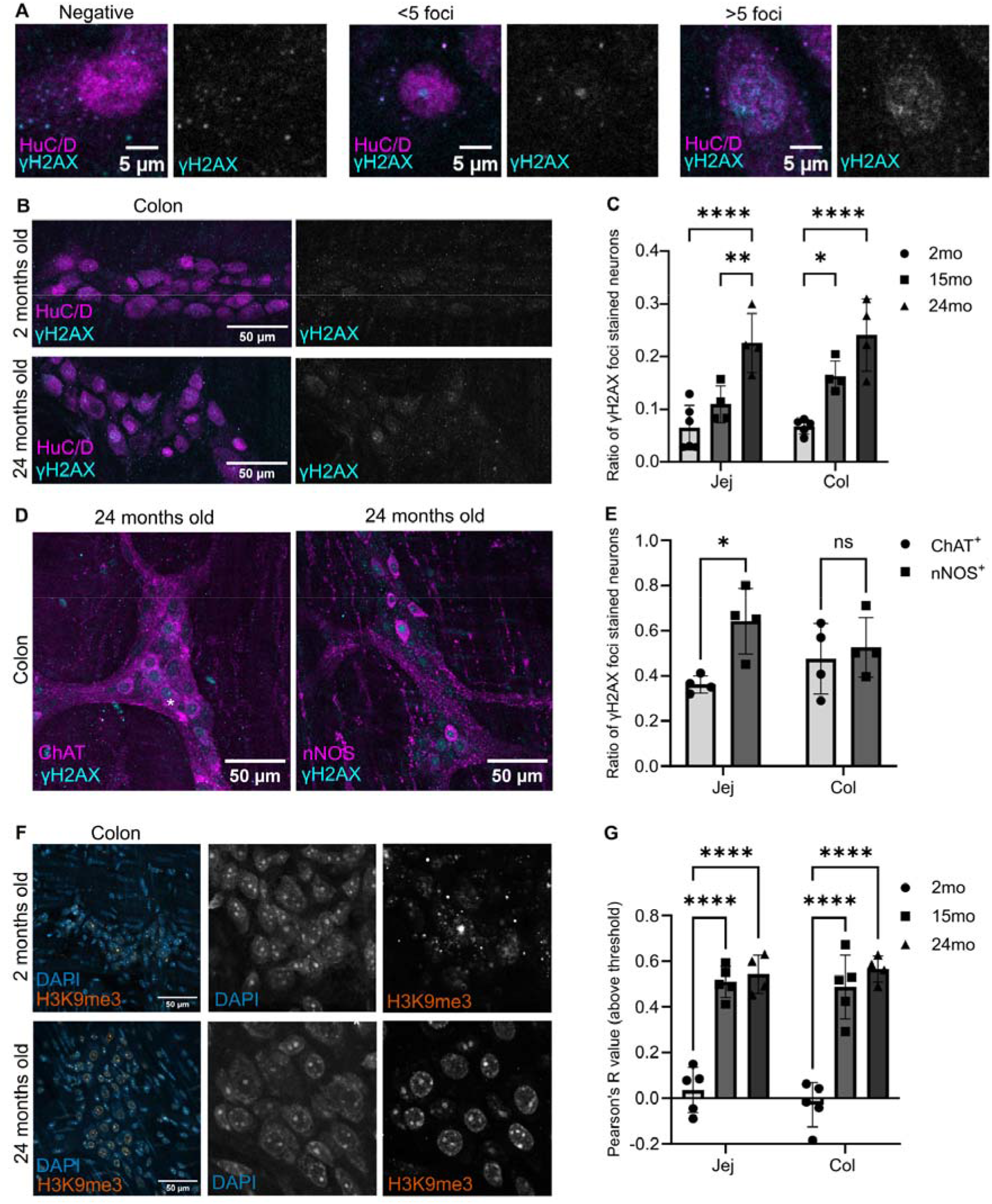
Enteric neurons accumulate DNA damage and epigenetic alterations with age. (A) Illustration of the scoring criteria for γH2AX positivity. From left to right: neuron without foci, neuron with <5 foci, and neuron with >5 foci (classified as γH2AX-positive). (B) Representative γH2AX and HuC/D co-stainings in young (top) and aged (bottom) colonic myenteric plexus. (C) Quantification of γH2AX-positive neurons in jejunum and colon across age groups. (D) Immunofluorescence images of aged myenteric ganglia stained for ChAT/HuC/D (left) and nNOS/HuC/D (right). (E) Quantification of γH2AX-positive nitrergic and cholinergic neurons in the jejunum and colon of 24-month-old mice. (F) Representative images of young (top) and aged (bottom) colonic myenteric ganglia immunolabeled for H3K9me3 and DAPI. (G) Pearson’s R quantification of H3K9me3-DAPI colocalization in 15- and 24-month-old jejunum and colon compared to 2-month samples. Every datapoint represents data from one animal.

Beyond genomic instability, we asked whether enteric neurons also undergo age-related epigenetic alterations, a second primary hallmark of aging. We assessed the formation of senescence-associated heterochromatin foci (SAHF) by staining for H3K9me3, a heterochromatin marker enriched in SAHF, together with the nuclear counterstain DAPI (Fig. 1F, Supplementary fig. 1B). Myenteric ganglia of aged mice showed increased H3K9me3-DAPI colocalization compared to those obtained from young mice, indicating enhanced chromatin condensation. Pearson’s R analysis confirmed a significant age-dependent increase in H3K9me3-DAPI colocalization in both jejunum and colon at 15 and 24 months (Fig. 1G). Notably, these epigenetic alterations were already fully present at 15 months, whereas DNA damage accumulation was more gradual and more prominent at 24 months, suggesting that epigenetic remodeling may precede the onset of more severe genomic instability during enteric neuronal aging.

### Enteric neuron loss and acquisition of a senescent phenotype with age

As mentioned earlier, aging has been linked to a progressive loss of enteric neurons within the ENS (12,29,30), alongside the emergence of features of cellular senescence (18). HuC/D staining showed a significantly reduced neuron density in the aged colon (24 vs 2 months), whereas neuron numbers in the jejunum were unchanged (Fig. 2A,B). Neuronal soma area increased modestly but significantly with age in both regions (Fig. 2C,D), consistent with cellular senescence (31). Staining for the cell-cycle inhibitor p21, a marker of senescence-associated arrest, was absent in young myenteric neurons but showed cytoplasmic signal in aged tissue, and at 24 months approximately 25-30% of neurons in both regions were p21-positive (Fig. 2E,F). Together, these findings indicate that aging enteric neurons are lost in the colon and acquire features of senescence, including soma enlargement and p21 expression, which may contribute to age-dependent ENS dysfunction.

**Figure 2.**
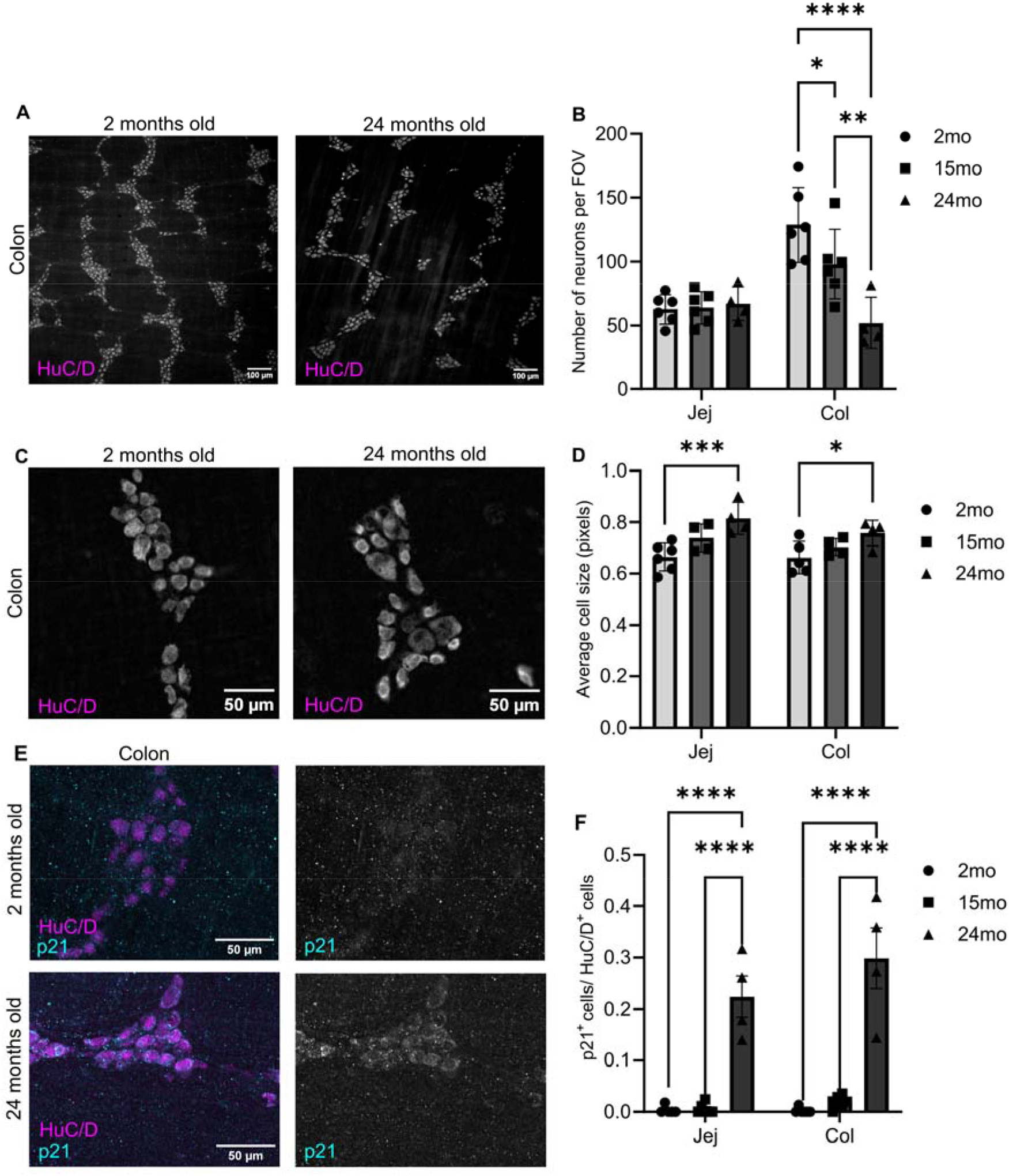
Enteric neuron loss and acquisition of a senescent phenotype with age. (A) Representative HuC/D immunostainings of colonic LMMP preparations from young (2 mo) and aged (24 mo) mice. (B) Quantification of HuC/D⁺ neurons per field of view in the jejunum and colon of 2-, 15-, and 24-month-old mice. (C) Representative HuC/D immunostainings of young (2 mo, left) and aged (24 mo, right) colonic LMMPs used for neuronal area analysis. (D) Quantification of neuronal area in the jejunum and colon across age groups. (E) Representative images of young (top) and aged (bottom) colonic myenteric ganglia immunolabeled for p21 and HuC/D. (F) Quantification of p21-positive neurons in the jejunum and colon across age groups. Every datapoint represents data from one animal.

### DSS-induced inflammation does not induce aging-associated senescence hallmarks in enteric neurons

To determine whether the aging-associated hallmarks reported above also occur under other forms of exogenous organismal stress, we examined the enteric neuron phenotype of 3 months old mice subjected to DSS-induced intestinal inflammation, a well-established model for colitis. γH2AX staining revealed a subtle increase in DNA damage in DSS-treated animals compared to healthy controls, visible both in enteric neurons and surrounding smooth muscle cells (Fig. 3A), which did not reach statistical significance (Fig. 3B). We next assessed the formation of SAHFs by evaluating the colocalization of H3K9me3 with DAPI. Representative images showed no apparent differences between control and tissues isolated from DSS-treated animals (Fig. 3C), and Pearson’s R analysis confirmed that H3K9me3-DAPI colocalization remained unchanged in both intestinal regions (Fig. 3D). Finally, we evaluated the expression of p21. Both healthy and myenteric plexus preparations from DSS-treated mice displayed minimal to no detectable p21 staining (Fig. 3E), and quantification confirmed near-zero levels of p21-positive neurons in both jejunum and colon across conditions (Fig. 3F). Together, the evidence that DNA damage accumulation, SAHF formation, and p21 induction do not arise in response to DSS-induced inflammation suggests that the aging hallmarks observed in aged mice (see earlier) are not universal signs of ENS stress but instead are specific to physiological aging.

**Figure 3.**
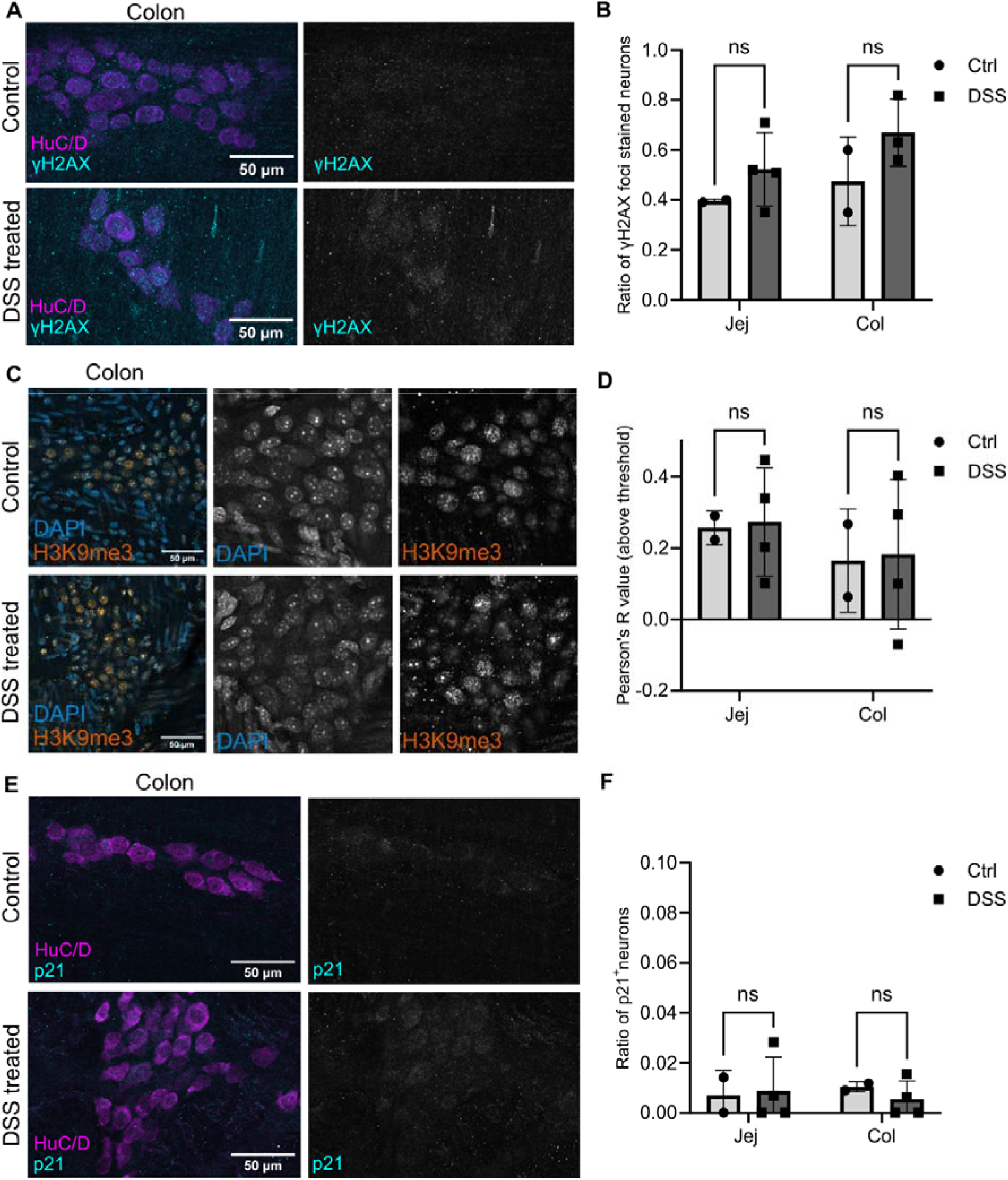
DSS-induced inflammation does not induce aging-associated senescence hallmarks in enteric neurons. (A) Representative γH2AX immunostainings of colonic LMMP preparations from healthy and DSS-treated mice. (B) Quantification of γH2AX-positive neurons in the jejunum and colon of healthy and DSS-treated mice. (C) Representative images of healthy (top) and DSS-treated (bottom) colonic myenteric ganglia stained for H3K9me3 and DAPI. (D) Quantification of H3K9me3-DAPI colocalization (Pearson’s R) in the jejunum and colon of healthy and DSS-treated mice. (E) Representative images of healthy (top) and DSS-treated (bottom) colonic myenteric ganglia immunolabeled for p21 and HuC/D. (F) Quantification of p21-positive neurons in the jejunum and colon of healthy and DSS-treated mice. Every datapoint represents data from one animal.

### Aging is associated with mitochondrial structural alterations and reduced metabolic activity in enteric neurons

To determine the effects of aging on mitochondrial structural integrity, we examined myenteric ganglion cross sections from young and aged mice using transmission electron microscopy (TEM). Quantification of mitochondrial morphology within enteric neuronal cell bodies and neurites revealed clear age-dependent changes (Fig. 4A, Supplementary fig. 2A). While mitochondria in young tissues appeared compact and uniformly shaped, those in aged samples were noticeably enlarged and displayed irregular outlines (Fig. 4A, white arrowheads). Morphometric analysis uncovered that mitochondrial surface area increased significantly with age (Fig. 4B), whereas mitochondrial circularity decreased (Fig. 4C), indicating a shift toward enlarged and structurally abnormal mitochondria. The observed morphological changes suggest the emergence of mitochondrial dysfunction, a central antagonistic hallmark, in the aging ENS.

**Figure 4.**
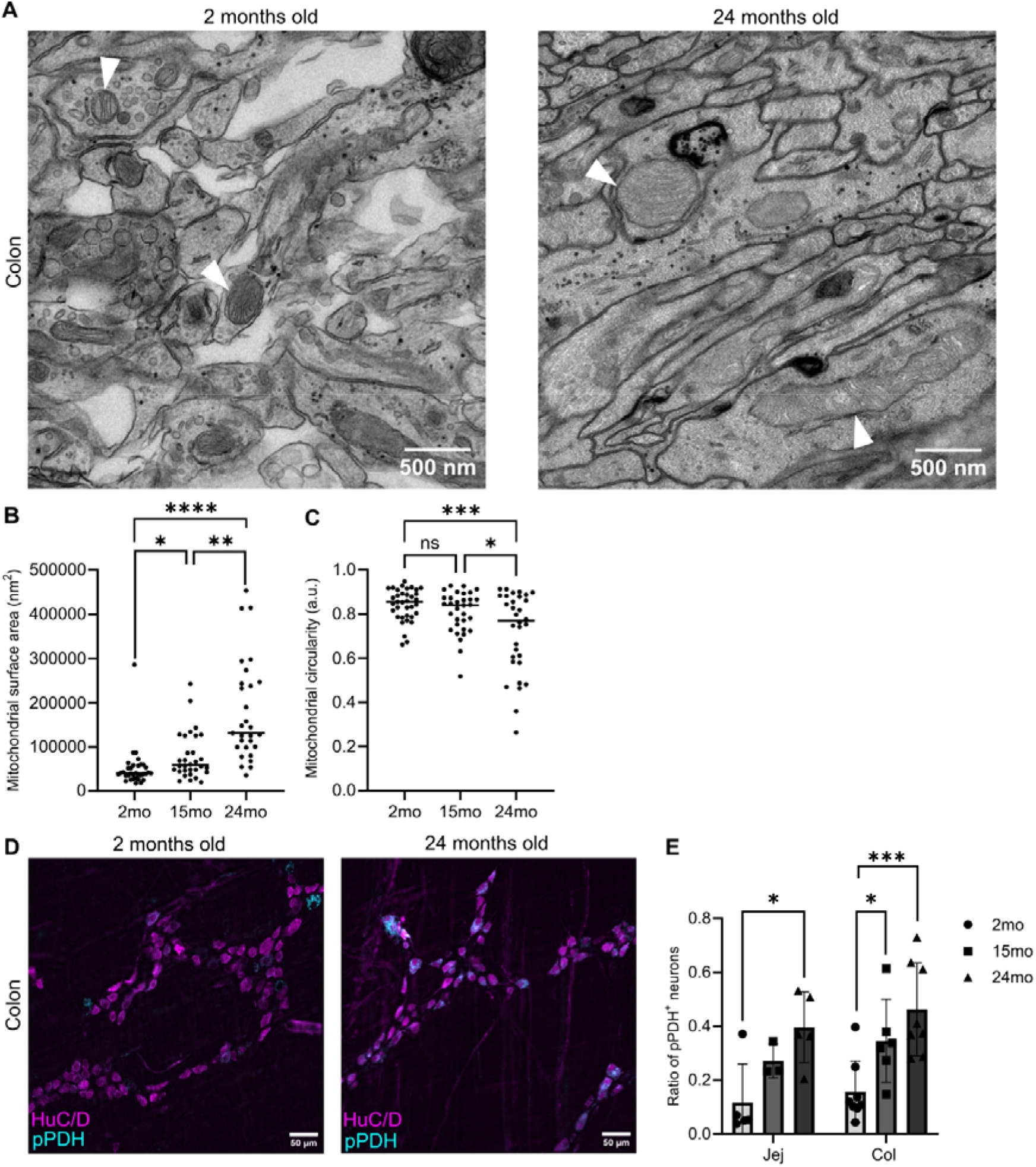
Aging is associated with mitochondrial structural alterations and reduced metabolic activity in enteric neurons. (A) Representative transmission electron microscopy images of colonic LMMP cross-sections from young (2 mo) and aged (24 mo) mice. White arrowheads indicate mitochondria. (B) Quantification of mitochondrial surface area. Each datapoint represents a single mitochondrion; mitochondria were pooled from 4 mice per group. (C) Quantification of mitochondrial circularity. Each datapoint represents a single mitochondrion; mitochondria were pooled from 4 mice per group. (D) Representative images of young (2 mo, left) and aged (24 mo, right) colonic LMMPs immunolabeled for pPDH and HuC/D. (E) Quantification of pPDH-positive neurons in the jejunum and colon of young and aged mice. Every datapoint represents data from one animal.

To determine whether the mitochondrial structural alterations are accompanied by functional changes in neuronal metabolism, we assessed the expression of phosphorylated pyruvate dehydrogenase (pPDH), a metabolic marker associated with reduced neuronal activity in both the central nervous system (CNS) and ENS (32,33). Immunostaining of colonic myenteric plexus preparations revealed markedly higher pPDH signal in aged enteric neurons compared to young controls (Fig. 4D, Supplementary fig. 2B). Quantification of the proportion of pPDH-positive neurons demonstrated a significant age-dependent increase in both the jejunum and colon (Fig. 4E), indicating that aging is accompanied by a shift toward a less active metabolic neuronal state. Collectively, the findings indicate that aging enteric neurons display both structural mitochondrial abnormalities and functional signatures consistent with reduced neuronal activity.

### Age-associated remodeling of enteric neuronal networks and divergent functional responses

Next, we asked whether aging alters enteric neuron function. To address this, we performed Ca²⁺ imaging on lmyenteric plexus preparations isolated from Wnt1-GCaMP3 mice. Electrical fiber tract stimulation (20 pulses, 300 μsec, 20 Hz) elicited increased Ca^2+^ responses in myenteric neurons of aged mice (Fig. 5A, B). Specifically, the Ca²⁺ transient amplitude of the response was significantly higher in aged animals compared to young controls (Fig. 5C). In contrast, the Ca²⁺ transient half-width was reduced in aged mice (Fig. 5D), indicating that aging alters both the magnitude and kinetics of electrically evoked neuronal network activity. We next examined receptor-mediated responses. Stimulation with the nicotinic agonist 1,1-dimethyl-4-phenylpiperazinium (DMPP) produced reduced responses in aged mice, with both Ca²⁺ transient amplitude and Ca²⁺ transient half-width significantly decreased compared to young animals (Fig. 5E–H). Similarly, adenosine triphosphate (ATP) stimulation resulted in lower Ca²⁺ transient amplitude and shorter Ca²⁺ transient half-width values in aged mice (Fig. 5I–L). Neurons were identified based on their characteristic GCaMP fluorescence pattern, including a large nuclear exclusion region and an oval-shaped neuronal cell body. Together, the results indicate that aging differentially affects neuronal responses, with enhanced responses to electrical stimulation but reduced responses to receptor-mediated activation.

**Figure 5.**
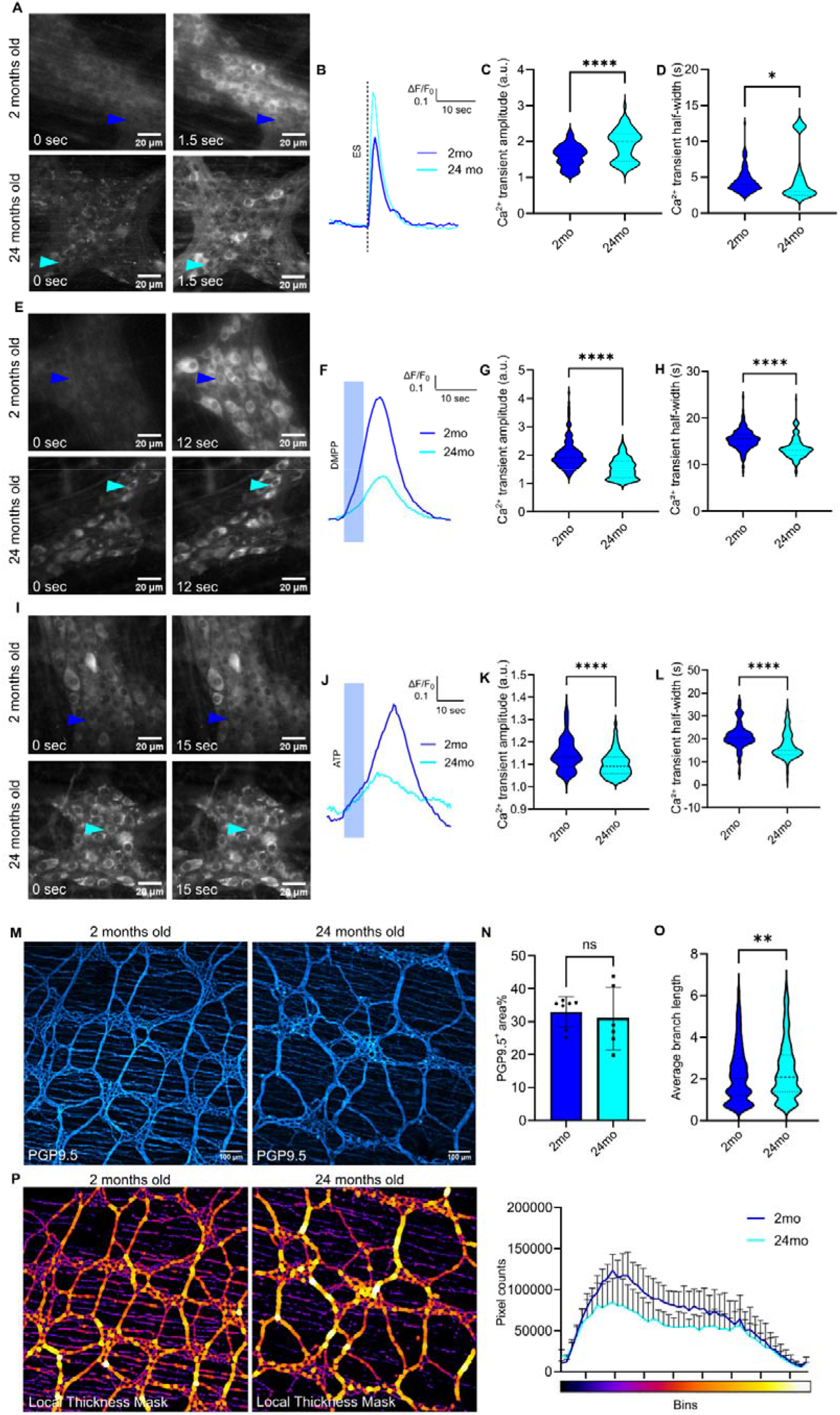
Age-associated remodeling of enteric neuronal networks leads to divergent functional responses. (A) Representative Ca²⁺ imaging snapshots of myenteric ganglia from young and aged Wnt1-GCaMP3 mice at rest (left) and during electrical train stimulation (20 Hz, 20 pulses) of a connected interganglionic fiber tract (right). Responding neurons are visualized by increased GCaMP fluorescence. (B) Example Ca²⁺ traces from individual neurons in young and aged mice following electrical stimulation. (C–D) Quantification of Ca²⁺ responses to electrical stimulation showing Ca²⁺ transient amplitude (C) and Ca²⁺ transient half-width (D) in young and aged mice. Each data point represents a single neuron; neurons were pooled from 3 mice per group. (E–H) Ca²⁺ responses to stimulation with the nicotinic agonist DMPP (10mM, 10sec). Representative traces (E–F) and quantification showing Ca²⁺ transient amplitude (G) and Ca²⁺ transient half-width (H) in young and aged mice. Each data point represents a single neuron; neurons were pooled from 3 mice per group. (I–L) Ca²⁺ responses following ATP stimulation (10mM, 10sec). Representative traces (I–J) and quantification demonstrating Ca²⁺ transient amplitude (K) and Ca²⁺ transient half-width (L) in young and aged mice. Each data point represents a single neuron; neurons were pooled from 3 mice per group. (M) Representative PGP9.5 immunostaining of neuronal processes in the colonic myenteric plexus of young and aged mice. (N) Quantification of total PGP9.5-positive area fraction per image (each data point represents one mouse). (O) Skeletonization-based network analysis showing neuronal fiber branch length in aged mice (each data point represents an individual branch derived from 7–8 mice per group). (P) Local thickness analysis of neuronal fiber bundles showing the distribution of bundle thicknesses in young and aged samples, revealing a highly significant age-related difference (p < 0.0001, Wilcoxon test).

The above findings suggest that age-related remodeling of the enteric neuronal network might contribute to the altered functional responses. To investigate potential structural changes, we analyzed myenteric plexus architecture using ubiquitin C-terminal hydrolase L1 (PGP9.5) immunostaining (Fig. 5M). Quantification of the total PGP9.5-positive fraction of the total area per image showed no significant difference between young and aged animals (Fig. 5N). However, skeletonization-based network analysis revealed a significant increase in fiber branch length in aged mice (Fig. 5O). In addition, local thickness analysis demonstrated a subtle but significant (p < 0.0001) shift in the distribution of fiber bundle width between young and aged samples (Fig. 5P). It should be noted that network analyses may be affected by differences in intramuscular nerve fiber content. As both age-related biological variation and technical variability during tissue dissection can influence this parameter, their relative contributions cannot be readily distinguished in the current study. Together, the results indicate that aging is associated with modest structural remodeling of the myenteric plexus, which may contribute to the divergent functional responses observed in Ca²⁺ imaging experiments.

### Age-associated hallmarks identified in mice are partially recapitulated in the human ENS

To determine whether the aging-associated changes observed in mice are also present in humans, we assessed several hallmark markers in enteric neurons from young and aged human gut samples. First, we examined DNA damage by co-staining for HuC/D and γH2AX (Fig. 6A). We also evaluated the SAHF marker H3K9me3 (Fig. 6B) and the neuronal inactivity marker pPDH in enteric neurons from young and aged individuals (Fig. 6C). Quantification of mean fluorescence intensity revealed a significant age-associated increase in neuronal γH2AX levels, indicating elevated DNA damage in aged enteric neurons (Fig. 6D). Similarly, H3K9me3 intensity was significantly increased with age (Fig. 6E). In contrast, pPDH levels were not significantly different between young and aged samples (Fig. 6F). In addition to phenotyping of aged enteric neurons, we investigated collagen deposition using second harmonic generation (SHG) imaging as a measure of tissue fibrosis and stiffness. This analysis was motivated by our mouse network analysis, which suggested that age-related remodeling of the ENS may be associated with changes in the surrounding extracellular matrix. Examination of human gut sections revealed prominent collagen accumulation between muscle fibers in both the longitudinal and circular muscle layer of aged tissues (Fig. 6G, H). Finally, we assessed the relationships between all measured parameters using a Pearson correlation matrix (Fig. 6I). Most parameters showed strong positive correlations with one another, suggesting that these aging-associated features occur in concert. In contrast, pPDH did not correlate significantly with any of the other measured parameters, consistent with the absence of age-related changes in its levels. In combination, these results indicate that aging of the human ENS is associated with increased DNA damage, enhanced senescence-associated chromatin remodeling, and increased collagen deposition, mirroring several of the age-related changes observed in mice.

**Figure 6.**
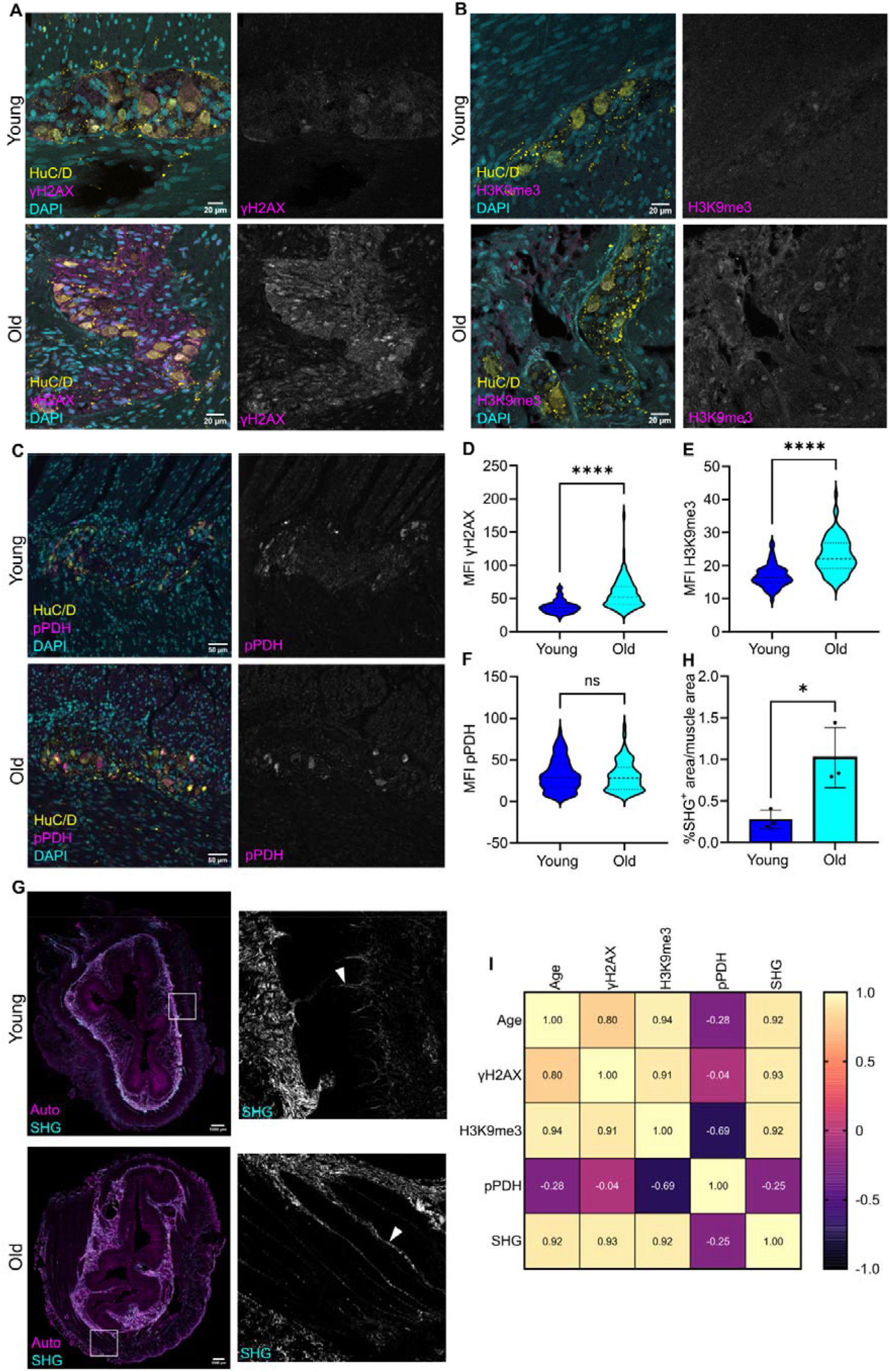
Age-associated hallmarks identified in mice are partially recapitulated in the human enteric nervous system. (A) Representative images of young and aged human gut sections immunolabeled for γH2AX and HuC/D. (B) Representative images of young and aged human enteric neurons stained for H3K9me3. (C) Representative images of young and aged human enteric neurons immunolabeled for pPDH and HuC/D. (D-F) Quantification of γH2AX (D), H3K9me3 (E), and pPDH (F) mean fluorescence intensity in HuC/D-positive enteric neurons from young and aged individuals. Each datapoint represents a single neuron; neurons were pooled from 3 individuals per group. (G) Representative second harmonic generation (SHG) images of collagen in young and aged human gut tissue. The white arrows indicate collagen deposition between the muscle layers. (H) Quantification of collagen content in the muscle layer based on SHG signal intensity. Every datapoint represents data from one individual. (I) Pearson correlation matrix showing relationships between neuronal aging markers and collagen content.

## DISCUSSION

In this study, we delineate the temporal emergence of several key molecular hallmarks of aging in enteric neurons of mice and humans. In mice, enteric neurons showed signs of genomic stress and epigenetic alteration with age, with γH2AX-positive neurons increasing gradually with age in both the jejunum and colon, and H3K9me3-DAPI colocalization, a sign of SAHF, emerging in full at 15 months. The DNA damage burden differed between neuronal subtypes in a region-specific manner, with nNOS-positive neurons in the jejunum carrying more γH2AX in aged samples than aged ChAT-positive neurons. Aged neurons also acquired features of cellular senescence beyond SAHF, including soma enlargement in both GI tract regions and visible p21 expression in roughly 25-30% of neurons at 24 months, along with a colon-restricted reduction in neuron number. Aging further affected mitochondria and metabolism, with ultrastructural analysis revealing enlarged, less circular mitochondria. Additionally, the proportion of pPDH-positive neurons increased in both regions, consistent with a less metabolically active neuronal state. Functionally, aging altered colonic neuronal activity in a stimulus-dependent manner, with enhanced Ca^2+^ responses to electrical stimulation but reduced nicotinergic and purinergic responses, together with a modest remodeling of the neuronal network architecture. Notably, DSS-induced colitis did not reproduce the DNA damage, SAHF, or p21 phenotypes seen with age, suggesting that these hallmarks are not general consequences of inflammatory stress but are specific to aging itself. Finally, in human tissue, enteric neurons showed an age-associated increase in γH2AX and H3K9me3 staining intensity, and aged muscle layers displayed increased collagen deposition, whereas, unlike in mice, pPDH did not change with age. Together, the data position DNA damage, chromatin remodeling, and neuronal senescence as core features of mammalian ENS aging, accompanied by mitochondrial, metabolic, and functional changes that so far have been demonstrated in mice.

Consistent with one of the fundamental hallmarks of aging, we observed an age-associated accumulation of DNA damage in both the jejunal and colonic ENS. This finding is in line with earlier observations by Jurk et al., who similarly reported increased γH2AX staining with age (18), supporting the notion that age-associated DNA damage signaling accumulates in enteric neurons. Such macromolecular damage accumulation is biologically plausible given that enteric neurons are terminally differentiated and long-lived (34,35), limiting the dilution of DNA lesions through cell division and favoring progressive buildup over the lifespan. While relatively few studies have investigated DNA damage specifically within the aging ENS, extensive evidence from other divisions/parts of the nervous system supports this concept. For example, γH2AX-positive nuclei increase from early life to old age in the mouse brain (36,37). Moreover, the age-associated accumulation of physical DNA lesions in cortical neurons was demonstrated by Mandavilli and Rao in 1996 (38), who reported that increased DNA damage did not inevitably trigger neuronal apoptosis, a finding that may explain why we observed substantial DNA damage accumulation in the jejunum despite the absence of overt neuronal loss. Thus, aging enteric neurons may similarly tolerate considerable levels of genomic damage before undergoing cell death.

DNA damage accumulation in enteric neurons was not uniform across neuronal populations. Specifically, we observed differences in susceptibility between cholinergic and nitrergic neuronal subtypes, indicating that neuronal vulnerability to aging-related genomic stress is heterogeneous. Similarly, Rutten et al. reported marked differences in the accumulation of DNA strand breaks between hippocampal granule cells, cerebellar granule cells and Purkinje neurons (39). Moreover, neuronal loss was only observed in Purkinje neurons, suggesting that DNA damage accumulation alone may be insufficient to drive neurodegeneration and that neuronal populations differ in their capacity to withstand genomic stress. The neuronal-specific differences in DNA damage accumulation and tolerance reported herein likely reflect subtype-specific disparities in: neuronal activity, which itself can induce DNA double-strand breaks (40); NO production, which has been linked to neuronal DNA strand breaks (41); or stress response/resilience mechanisms, including DNA repair capacity (42). Moreover, given the regional differences in ENS cell composition, physiological function and metabolic demands (43–45), these factors likely influence the susceptibility of particular gut segments (e.g., colon and jejunum) to aging-related stressors.

In addition to increased DNA damage, we observed a significant increase in the correlation between DAPI and H3K9me3 staining from 15 months onward, indicative of age-associated changes in chromatin organization. Notably, the peak DAPI-H3K9me3 colocalization emerged before the highest levels of DNA damage accumulation were observed, suggesting that chromatin remodeling represents an early event in enteric neuronal aging. This staining pattern is also consistent with the formation of SAHF, a hallmark of cellular senescence originally described in fibroblasts by Narita et al. (46). While SAHF have been reported across a range of aging tissues since, including post-mitotic cell populations (47), the work herein represents the first identification of SAHF in the ENS. Since heterochromatin and the DNA damage response are mechanistically intertwined (47,48), reorganization of H3K9me3-marked chromatin could alter the efficiency of DNA damage signalling and repair, providing a plausible route by which early chromatin changes contribute to later damage accumulation. Mechanistically, their formation is thought to result from the spatial reorganization of pre-existing H3K9me3-marked heterochromatin domains rather than an increase in global H3K9me3 abundance (49). Collectively, our findings suggest that age-associated chromatin reorganization occurs early within the ENS and may constitute a previously unrecognized driver of enteric neuronal aging.

Although the functional implications of increased H3K9me3 remodelling in aging enteric neurons remain unclear, accumulating evidence suggests that alterations in heterochromatin organization can significantly influence neuronal function (50,51). For example, Snigdha et al. demonstrated that pharmacological reduction of H3K9me3 through inhibition of SUV39H1 improved memory performance and synaptic function in aged mice, suggesting that excessive heterochromatinization may contribute to age-associated neuronal dysfunction in the brain (50). At the same time, changes in H3K9me3 appear to be highly cell type specific, with heterochromatin loss and reduced H3K9me3 levels detected in aged excitatory cortical neurons being largely absent from inhibitory neurons and glial cells (51). These observations caution against assuming a uniform aging-associated chromatin response across neuronal populations and may indicate that enteric neurons engage distinct epigenetic aging programs compared with central neurons.

Alongside DNA damage and chromatin remodeling, aged enteric neurons displayed two additional features of cellular senescence: progressive soma hypertrophy and p21 expression. Neuronal soma size increased gradually with age; and by 24 months, approximately 25-30% of neurons were p21-positive, whereas p21-positive neurons were not detected at the earlier time points. Cell enlargement is a long-recognized hallmark of senescence, and recent work suggests that it can be both a consequence and a cause of permanent cell cycle exit (31). Moreover, cellular hypertrophy has been reported in other aged post-mitotic cells with senescence markers, such as cardiomyocytes (reviewed in (52)).

However, we cannot rule out the possibility that selective loss of enteric neuron subtypes with smaller cell soma, contributes to shifts in average neuronal soma size (53,54). Therefore, while the observed hypertrophy is consistent with a senescence-associated phenotype, part of the effect may also reflect age-dependent alterations in neuronal subtype composition. The p21 data agree with the original description of a p21-dependent, senescence-like phenotype in post-mitotic neurons by Jurk et al., who found multiple senescence markers co-occurring in the same neurons of old but not young mice (18), including central neurons and neurons of the myenteric plexus (52). In that model, persistent DNA damage, driven in part by dysfunctional telomeres, activates the ataxia-telangiectasia mutated (ATM)/ATM and Rad3-related (ATR)-Chk-p53 axis and induces p21, which inhibits cyclin-dependent kinase (CDK) activity and enforces cell cycle arrest (52). This paradigm is consistent with our observation that p21 appears at a time when DNA damage burden is high.

The mitochondrial and metabolic changes we observed in aged enteric neurons are relevant in the context of our senescence findings. In line with our observation that mitochondria were larger and more irregular in shape in aged enteric neurons, comparable alterations have been described in other aged neuronal populations. Specifically, oversized synaptic mitochondria were seen in old monkey cerebellum biopsies and found to exhibit reduced cytochrome oxidase-based metabolic competence (55); aged rat hippocampal synapses were shown to contain more structurally damaged mitochondria (56); and enlarged but structurally simpler mitochondria were reported in axons of aged rat hypoglossal motor neurons (57). Paired with the increased proportion of pPDH-positive neurons, our work suggests that enlarged, irregular mitochondria in aged enteric neurons may reflect functional impairment rather than an adaptive change, consistent with the role of mitochondrial dysfunction in senescence (52). Altogether, the current results support the idea that enteric neurons mount a senescence-like response with age. Our data from an independent collection of human samples add independent evidence of post-mitotic senescence, specifically in the ENS.

Functionally, aged enteric neurons displayed enhanced electrically evoked Ca²⁺ responses and reduced responses to agonist stimulation. Given the prominent role of nicotinergic receptors in mediating excitatory enteric neurotransmission in the colonic ENS (28), the apparent paradoxical finding of reduced DMPP-evoked responses alongside enhanced responses to network activation suggests compensatory remodeling, altered connectivity and functional rewiring of aged enteric neural circuits. Together with the significant enteric neuron loss, these findings support the idea that the ENS has considerable functional reserve and can maintain output despite age-related cellular changes (58). Impaired activity of individual enteric neurons may also be linked to the senescence-like phenotype, since Ca²⁺ handling depends on mitochondrial buffering and ATP supply, and the enlarged, irregular mitochondria and increased pPDH in aged neurons suggest that these elements are altered. Additionally, the structural remodeling of the neuronal network, with longer fiber branches but unchanged overall PGP9.5 area, is in line with the swollen neuronal processes reported in aged mouse colon (59) and with the morphological changes that accompany senescence, such as soma enlargement. Lastly, cytoplasmic p21 has been shown to regulate neurite remodeling through inhibition of Rho-kinase (60), which suggests a possible link between the p21 expression we observed and the altered network architecture. Together, the current evidence suggests that aging-associated senescence-like changes in enteric neurons extend from the molecular level to cellular function and network organization, although it remains to be determined whether the altered responses are compensatory.

Preliminary studies conducted on human gut samples found that aged human enteric neurons exhibit higher γH2AX and H3K9me3 intensity, a finding consistent with our mouse data. While the readouts were distinct, i.e., H3K9me3 was quantified as mean intensity in human tissue and as DAPI colocalization in mice, both results point to an age-associated change in heterochromatin. Notably, in contrast to mice, pPDH did not differ significantly with age in the human samples. This apparent discrepancy could stem from several reasons, including the small sample size, low statistical power, greater heterogeneity among humans, species differences, or the lability of phospho-epitopes in surgical tissue, meaning we cannot draw any major conclusions from the absence of an effect. Aged human muscle layers also showed more collagen than young mice, an observation that is in line with earlier reports of higher total collagen in the submucosa and muscularis externa of the elderly human colon (20). While the neuronal network remodeling and extracellular matrix changes were observed in mice and humans, respectively, both findings point toward age-associated structural remodeling of the GI neuromuscular environment. However, the present study cannot establish a direct relationship between these processes. Overall, our early investigations suggest that some features of enteric neuronal aging, namely DNA damage, chromatin remodeling and fibrotic tissue changes, may be conserved between mouse and human. Confirmation in larger, well-characterized cohorts is needed.

Several limitations of the work herein should be pointed out. First, the study design is cross-sectional, so we cannot firmly establish the order of events or causality between chromatin remodeling, DNA damage, senescence-like changes and altered function. Second, as already mentioned, the human cohort was very small (three individuals per group) and only covered a subset of markers. Third, the Ca²⁺ imaging was performed in colonic preparations from male mice only, and the DSS cohort was also relatively small. Consequently, the results of these experiments should be interpreted with appropriate caution. Despite these limitations, our data suggest that aging enteric neurons develop a coordinated, senescence-like phenotype, involving DNA damage, chromatin reorganization, p21 expression, mitochondrial alterations and altered functional responses, which differs from what is induced by inflammatory stress and may be at least partly conserved in humans. The findings presented here lay a foundation for future studies using larger human cohorts, subtype-resolved and longitudinal analyses, and functional manipulation of senescent neurons to determine whether these changes drive age-related GI dysfunction and whether they can be targeted therapeutically.

## AUTHOR CONTRIBUTIONS

Lobke Mombeek: Conceptualization, Data curation, Formal analysis, Funding acquisition, Investigation, Methodology, Project administration, Supervision, Validation, Visualization, Writing – original draft, Writing – review & editing. Anton Brosens: Investigation (p21 staining and γH2AX staining in neuronal subtypes), Formal analysis. Reindert Jehoul: Investigation (DSS-induced stressor experiments). Anita Leembruggen: Investigation (calcium imaging experiments). Peter Neckel: Resources (provision of human samples). Marlene Hao: Investigation (calcium imaging experiments), Methodology, Resources. Werend Boesmans: Conceptualization, Funding acquisition, Supervision, Writing – review & editing. David Wilson III: Conceptualization, Funding acquisition, Supervision, Writing – review & editing.

## REFERENCES

1. Furness JB. The enteric nervous system and neurogastroenterology. Nat Rev Gastroenterol Hepatol 2012 95 [Internet]. 2012 Mar 6 [cited 2024 Feb 8];9(5):286–94. Available from: https://www.nature.com/articles/nrgastro.2012.32

2. Costa M, Brookes SJH, Hennig GW. Anatomy and physiology of the enteric nervous system. Gut [Internet]. 2000 [cited 2026 Oct 22];47 Suppl 4(Suppl 4). Available from: https://pubmed.ncbi.nlm.nih.gov/11076898/

3. Sharkey KA, Mawe GM. The enteric nervous system. Physiol Rev [Internet]. 2023 Apr 1 [cited 2026 Oct 22];103(2):1487–564. Available from: https://pubmed.ncbi.nlm.nih.gov/36521049/

4. Stavely R, Ott LC, Rashidi N, Sakkal S, Nurgali K. The Oxidative Stress and Nervous Distress Connection in Gastrointestinal Disorders. Biomolecules [Internet]. 2023 Nov 1 [cited 2026 Oct 22];13(11). Available from: https://pubmed.ncbi.nlm.nih.gov/38002268/

5. Chow AK, Gulbransen BD. Potential roles of enteric glia in bridging neuroimmune communication in the gut. Am J Physiol - Gastrointest Liver Physiol [Internet]. 2016 [cited 2026 Oct 22];312(2):G145. Available from: https://pmc.ncbi.nlm.nih.gov/articles/PMC5338608/

6. Bellitto V, Tomassoni D, Martinelli I, Nittari G, Tayebati SK. Obesity-Related Oxidative Stress and Antioxidant Properties of Natural Compounds in the Enteric Nervous System: A Literature Overview. Antioxidants (Basel, Switzerland) [Internet]. 2026 Jan 1 [cited 2026 Oct 22];15(1). Available from: https://pubmed.ncbi.nlm.nih.gov/41596140/

7. 7. Mazzuoli-Weber G, Elfers K. Modulation of the microbiota-enteric nervous system axis by environmental factors: from adaptive plasticity to maladaptive neuroenteric remodeling. Front Neurosci [Internet]. 2026 [cited 2026 Oct 22];20. Available from: https://pubmed.ncbi.nlm.nih.gov/42819344/

8. 8. Sanchini G, Tevosian M, Cardilli A, Bou Abboud L, Uesaka T, Rosshart SP, et al. Environmental cues shape mucosal enteric glia networks independent of lamina propria innervation. npj Biofilms Microbiomes 2026 [Internet]. 2026 Oct 17 [cited 2026 Oct 22]; Available from: https://www.nature.com/articles/s41522-026-01175-5

9. López-Otín C, Blasco MA, Partridge L, Serrano M, Kroemer G. Hallmarks of aging: An expanding universe. Cell [Internet]. 2023 Jan 19 [cited 2026 Oct 22];186(2):243–78. Available from: https://pubmed.ncbi.nlm.nih.gov/36599349/

10. Schaum N, Lehallier B, Hahn O, Pálovics R, Hosseinzadeh S, Lee SE, et al. Ageing hallmarks exhibit organ-specific temporal signatures. Nat 2020 5837817 [Internet]. 2020 Jul 15 [cited 2026 Oct 22];583(7817):596–602. Available from: https://www.nature.com/articles/s41586-020-2499-y

11. Oh HSH, Rutledge J, Nachun D, Pálovics R, Abiose O, Moran-Losada P, et al. Organ aging signatures in the plasma proteome track health and disease. Nat 2023 6247990 [Internet]. 2023 Dec 6 [cited 2026 Oct 22];624(7990):164–72. Available from: https://www.nature.com/articles/s41586-023-06802-1

12. Abalo R, Rivera AJ, Vera G, Martín MI. Ileal myenteric plexus in aged guinea-pigs: loss of structure and calretinin-immunoreactive neurones. Neurogastroenterol Motil [Internet]. 2005 Feb [cited 2026 Oct 22];17(1):123–32. Available from: https://pubmed.ncbi.nlm.nih.gov/15670272/

13. Phillips RJ, Hudson CN, Powley TL. Sympathetic axonopathies and hyperinnervation in the small intestine smooth muscle of aged Fischer 344 rats. Auton Neurosci Basic Clin [Internet]. 2013 [cited 2026 Oct 22];179(1–2):108–21. Available from: https://pubmed.ncbi.nlm.nih.gov/24104187/

14. Nguyen TT, Baumann P, Tüscher O, Schick S, Endres K. The Aging Enteric Nervous System. Int J Mol Sci [Internet]. 2023 Jun 1 [cited 2026 Oct 22];24(11). Available from: https://pubmed.ncbi.nlm.nih.gov/37298421/

15. Deb B, Prichard DO, Bharucha AE. Constipation and Fecal Incontinence in the Elderly. Curr Gastroenterol Rep [Internet]. 2020 Nov 1 [cited 2026 Oct 22];22(11). Available from: https://pubmed.ncbi.nlm.nih.gov/32839874/

16. Wade PR. Aging and neural control of the GI tract. I. Age-related changes in the enteric nervous system. Am J Physiol Gastrointest Liver Physiol [Internet]. 2002 [cited 2026 Oct 22];283(3). Available from: https://pubmed.ncbi.nlm.nih.gov/12181159/

17. Szymaszkiewicz A, Szymaszkiewicz K, Fichna J, Zielińska M. The age-related alerations in the enteric nervous system and their impact on peristalsis of the gastrointestinal tract. Postepy Biochem [Internet]. 2021 Jan 13 [cited 2026 Oct 22];67(1):34–43. Available from: https://pubmed.ncbi.nlm.nih.gov/34378903/

18. Jurk D, Wang C, Miwa S, Maddick M, Korolchuk V, Tsolou A, et al. Postmitotic neurons develop a p21-dependent senescence-like phenotype driven by a DNA damage response. Aging Cell [Internet]. 2012 Dec [cited 2024 May 28];11(6):996. Available from: /pmc/articles/PMC3533793/

19. 19. Palmer A, Epton S, Crawley E, Straface M, Gammon L, Edgar MM, et al. Expression of p16 Within Myenteric Neurons of the Aged Colon: A Potential Marker of Declining Function. Front Neurosci [Internet]. 2021 Oct 7 [cited 2026 Oct 22];15. Available from: https://pubmed.ncbi.nlm.nih.gov/34690683/

20. Baidoo N, Crawley E, Knowles CH, Sanger GJ, Belai A. Total collagen content and distribution is increased in human colon during advancing age. PLoS One [Internet]. 2022 Jun 1 [cited 2026 Oct 22];17(6):e0269689. Available from: https://journals.plos.org/plosone/article?id=10.1371/journal.pone.0269689

21. De Schepper S, Verheijden S, Aguilera-Lizarraga J, Viola MF, Boesmans W, Stakenborg N, et al. Self-Maintaining Gut Macrophages Are Essential for Intestinal Homeostasis. Cell [Internet]. 2018 Oct 4 [cited 2026 Oct 22];175(2):400–415.e13. Available from: https://pubmed.ncbi.nlm.nih.gov/30173915/

22. Schindelin J, Arganda-Carreras I, Frise E, Kaynig V, Longair M, Pietzsch T, et al. Fiji: an open-source platform for biological-image analysis. Nat Methods 2012 97 [Internet]. 2012 Jun 28 [cited 2026 Oct 27];9(7):676–82. Available from: https://www.nature.com/articles/nmeth.2019

23. Danielian PS, Muccino D, Rowitch DH, Michael SK, McMahon AP. Modification of gene activity in mouse embryos in utero by a tamoxifen-inducible form of Cre recombinase. Curr Biol [Internet]. 1998 Dec 3 [cited 2026 Oct 22];8(24):1323–6. Available from: https://pubmed.ncbi.nlm.nih.gov/9843687/

24. Zariwala HA, Borghuis BG, Hoogland TM, Madisen L, Tian L, de Zeeuw CI, et al. A Cre-dependent GCaMP3 reporter mouse for neuronal imaging in vivo. J Neurosci [Internet]. 2012 Feb 29 [cited 2026 Oct 22];32(9):3131–41. Available from: https://pubmed.ncbi.nlm.nih.gov/22378886/

25. 25. Balasuriya GK, Nugapitiya SS, Hill-Yardin EL, Bornstein JC. Nitric Oxide Regulates Estrus Cycle Dependent Colonic Motility in Mice. Front Neurosci [Internet]. 2021 Sep 29 [cited 2026 Oct 22];15. Available from: https://pubmed.ncbi.nlm.nih.gov/34658750/

26. Vanden Berghe P, Kenyon JL, Smith TK. Mitochondrial Ca2+ Uptake Regulates the Excitability of Myenteric Neurons. J Neurosci [Internet]. 2002 Aug 15 [cited 2026 Oct 27];22(16):6962–71. Available from: https://www.jneurosci.org/content/22/16/6962

27. Leembruggen AJL, Yildiz GS, Hardee JP, Stamp LA, Bornstein JC, Hao MM. Plasticity of enteric neurotransmission varies during day-night cycles and with feeding state. Am J Physiol Gastrointest Liver Physiol [Internet]. 2025 Feb 1 [cited 2026 Nov 10];328(2):G145–51. Available from: https://pubmed.ncbi.nlm.nih.gov/39763332/

28. Li Z, Boesmans W, Kazwiny Y, Hao MM, Berghe P Vanden. Simultaneous whole-cell patch-clamp and calcium imaging on myenteric neurons. Am J Physiol - Gastrointest Liver Physiol. 2022 Oct 1;323(4):G341–7.

29. Bernard CE, Gibbons SJ, Gomez-Pinilla PJ, Lurken MS, Schmalz PF, Roeder JL, et al. Effect of Age on the Enteric Nervous System of the Human Colon. Neurogastroenterol Motil [Internet]. 2009 Jul [cited 2026 Oct 22];21(7):746. Available from: https://pmc.ncbi.nlm.nih.gov/articles/PMC2776702/

30. Phillips RJ, Kieffer EJ, Powley TL. Loss of glia and neurons in the myenteric plexus of the aged Fischer 344 rat. Anat Embryol (Berl). 2004 Nov;209(1):19–30.

31. Manohar S, Neurohr GE. Too big not to fail: emerging evidence for size-induced senescence. FEBS J [Internet]. 2024 Jun 1 [cited 2026 Oct 22];291(11):2291–305. Available from: https://pubmed.ncbi.nlm.nih.gov/37986656/

32. Yang D, Wang Y, Qi T, Zhang X, Shen L, Ma J, et al. Phosphorylation of pyruvate dehydrogenase inversely associates with neuronal activity. Neuron [Internet]. 2024 Mar 20 [cited 2026 Oct 22];112(6):959–971.e8. Available from: https://pubmed.ncbi.nlm.nih.gov/38266644/

33. Wemelle E, McClain JL, Gulbransen BD. Phosphorylation of Pyruvate Dehydrogenase as a Marker of Neuronal Inactivity in the Enteric Nervous System. Neurogastroenterol Motil [Internet]. 2025 Nov 1 [cited 2026 Oct 22];37(11):e70135. Available from: https://pmc.ncbi.nlm.nih.gov/articles/PMC12534579/

34. Virtanen H, Garton DR, Andressoo JO. Myenteric Neurons Do Not Replicate in Small Intestine Under Normal Physiological Conditions in Adult Mouse. CMGH [Internet]. 2022 Jan 1 [cited 2026 Oct 22];14(1):27–34. Available from: https://pubmed.ncbi.nlm.nih.gov/35421596/

35. Pham TD, Gershon MD, Rothman TP. Time of origin of neurons in the murine enteric nervous system: Sequence in relation to phenotype. J Comp Neurol [Internet]. 1991 Dec 22 [cited 2026 Oct 27];314(4):789–98. Available from: /doi/pdf/10.1002/cne.903140411

36. Barral S, Beltramo R, Salio C, Aimar P, Lossi L, Merighi A. Phosphorylation of histone H2AX in the mouse brain from development to senescence. Int J Mol Sci [Internet]. 2014 Jan 21 [cited 2026 Oct 22];15(1):1554–73. Available from: https://pubmed.ncbi.nlm.nih.gov/24451138/

37. Merighi A, Gionchiglia N, Granato A, Lossi L. The Phosphorylated Form of the Histone H2AX (γH2AX) in the Brain from Embryonic Life to Old Age. Molecules [Internet]. 2021 Dec 1 [cited 2026 Oct 22];26(23). Available from: https://pubmed.ncbi.nlm.nih.gov/34885784/

38. Mandavilli BS, Rao KS. Accumulation of DNA damage in aging neurons occurs through a mechanism other than apoptosis. J Neurochem [Internet]. 1996 [cited 2026 Oct 22];67(4):1559–65. Available from: https://pubmed.ncbi.nlm.nih.gov/8858940/

39. Rutten BPF, Schmitz C, Gerlach OHH, Oyen HM, de Mesquita EB, Steinbusch HWM, et al. The aging brain: Accumulation of DNA damage or neuron loss? Neurobiol Aging [Internet]. 2007 Jan [cited 2026 Oct 22];28(1):91–8. Available from: https://pubmed.ncbi.nlm.nih.gov/16338029/

40. Suberbielle E, Sanchez PE, Kravitz A V., Wang X, Ho K, Eilertson K, et al. Physiologic brain activity causes DNA double-strand breaks in neurons, with exacerbation by amyloid-β. Nat Neurosci [Internet]. 2013 May [cited 2026 Oct 22];16(5):613–21. Available from: https://pubmed.ncbi.nlm.nih.gov/23525040/

41. Pieper AA, Blackshaw S, Clements EE, Brat DJ, Krug DK, White AJ, et al. Poly(ADP-ribosyl)ation basally activated by DNA strand breaks reflects glutamate-nitric oxide neurotransmission. Proc Natl Acad Sci U S A [Internet]. 2000 Feb 15 [cited 2026 Oct 22];97(4):1845–50. Available from: https://www.pnas.org/doi/pdf/10.1073/pnas.97.4.1845

42. 42. Wang X, Michaelis EK. Selective neuronal vulnerability to oxidative stress in the brain. Front Aging Neurosci [Internet]. 2010 [cited 2026 Oct 22];2(MAR). Available from: https://pubmed.ncbi.nlm.nih.gov/20552050/

43. Hamnett R, Dershowitz LB, Sampathkumar V, Wang Z, Gomez-Frittelli J, De Andrade V, et al. Regional cytoarchitecture of the adult and developing mouse enteric nervous system. Curr Biol [Internet]. 2022 Oct 24 [cited 2026 Oct 22];32(20):4483–4492.e5. Available from: https://pubmed.ncbi.nlm.nih.gov/36070775/

44. Obata Y, Castaño Á, Boeing S, Bon-Frauches AC, Fung C, Fallesen T, et al. Neuronal programming by microbiota regulates intestinal physiology. Nat 2020 5787794 [Internet]. 2020 Feb 5 [cited 2026 Oct 27];578(7794):284–9. Available from: https://www.nature.com/articles/s41586-020-1975-8

45. Millett CJ, Shaver JJ, Bracken B, Jones SJ, Lovelett RJ, Rubinow DA, et al. In vivo transcriptomic, functional, circuit-based, and translational analyses of enteric neurons. Cell [Internet]. 2025 Dec 24 [cited 2026 Oct 27];188(26):7547–7570.e45. Available from: https://pubmed.ncbi.nlm.nih.gov/41406962/

46. Narita M, Nũnez S, Heard E, Narita M, Lin AW, Hearn SA, et al. Rb-mediated heterochromatin formation and silencing of E2F target genes during cellular senescence. Cell [Internet]. 2003 Jun 13 [cited 2026 Oct 22];113(6):703–16. Available from: https://pubmed.ncbi.nlm.nih.gov/12809602/

47. Cann KL, Dellaire G. Heterochromatin and the DNA damage response: the need to relax. Biochem Cell Biol [Internet]. 2011 Feb [cited 2026 Oct 22];89(1):45–60. Available from: https://pubmed.ncbi.nlm.nih.gov/21326362/

48. Goodarzi AA, Noon AT, Jeggo PA. The impact of heterochromatin on DSB repair. Biochem Soc Trans [Internet]. 2009 [cited 2026 Oct 22];37(Pt 3):569–76. Available from: https://pubmed.ncbi.nlm.nih.gov/19442252/

49. Chandra T, Kirschner K, Thuret JY, Pope BD, Ryba T, Newman S, et al. Independence of Repressive Histone Marks and Chromatin Compaction during Senescent Heterochromatic Layer Formation. Mol Cell [Internet]. 2012 Jul 27 [cited 2026 Oct 22];47(2):203–14. Available from: https://pubmed.ncbi.nlm.nih.gov/22795131/

50. Snigdha S, Aleph Prieto G, Petrosyan A, Loertscher BM, Dieskau AP, Overman LE, et al. H3K9me3 Inhibition Improves Memory, Promotes Spine Formation, and Increases BDNF Levels in the Aged Hippocampus. J Neurosci [Internet]. 2016 Mar 23 [cited 2026 Oct 22];36(12):3611–22. Available from: https://pubmed.ncbi.nlm.nih.gov/27013689/

51. Zhang Y, Amaral ML, Zhu C, Grieco SF, Hou X, Lin L, et al. Single-cell epigenome analysis reveals age-associated decay of heterochromatin domains in excitatory neurons in the mouse brain. Cell Res 2022 3211 [Internet]. 2022 Oct 7 [cited 2026 Oct 22];32(11):1008–21. Available from: https://www.nature.com/articles/s41422-022-00719-6

52. Von Zglinicki T, Wan T, Miwa S. Senescence in Post-Mitotic Cells: A Driver of Aging? Antioxid Redox Signal [Internet]. 2021 Feb 1 [cited 2026 Oct 22];34(4):308–23. Available from: https://pubmed.ncbi.nlm.nih.gov/32164429/

53. Qu ZD, Thacker M, Castelucci P, Bagyánszki M, Epstein ML, Furness JB. Immunohistochemical analysis of neuron types in the mouse small intestine. Cell Tissue Res 2008 3342 [Internet]. 2008 Oct 15 [cited 2026 Nov 10];334(2):147–61. Available from: https://link.springer.com/article/10.1007/s00441-008-0684-7

54. Nurgali K, Stebbing MJ, Furness JB. Correlation of electrophysiological and morphological characteristics of enteric neurons in the mouse colon. J Comp Neurol [Internet]. 2004 Jan 1 [cited 2026 Nov 10];468(1):112–24. Available from: /doi/pdf/10.1002/cne.10948

55. Bertoni-Freddari C, Balietti M, Giorgetti B, Grossi Y, Casoli T, Di Stefano G, et al. Selective decline of the metabolic competence of oversized synaptic mitochondria in the old monkey cerebellum. Rejuvenation Res [Internet]. 2008 Apr 1 [cited 2026 Oct 22];11(2):387–91. Available from: https://pubmed.ncbi.nlm.nih.gov/18328003/

56. Rybka V, Suzuki YJ, Gavrish AS, Dibrova VA, Gychka SG, Shults N V. Transmission Electron Microscopy Study of Mitochondria in Aging Brain Synapses. Antioxidants (Basel, Switzerland) [Internet]. 2019 Jun 1 [cited 2026 Oct 22];8(6). Available from: https://pubmed.ncbi.nlm.nih.gov/31212589/

57. 57. Searles MJ, Dasgupta D, Christensen TA, Sieck GC, Fogarty MJ. Age-related disruptions in ubiquitination and mitochondrial proteostasis are associated with neuronal compartment-specific mitochondrial changes. Exp Gerontol [Internet]. 2026 Oct 15 [cited 2026 Oct 22];224. Available from: https://pubmed.ncbi.nlm.nih.gov/42685870/

58. Saffrey MJ. Cellular changes in the enteric nervous system during ageing. Dev Biol [Internet]. 2013 Oct 1 [cited 2026 Oct 22];382(1):344–55. Available from: https://pubmed.ncbi.nlm.nih.gov/23537898/

59. Gamage PP, Ranson RN, Patel BA, Yeoman MS, Saffrey MJ. Myenteric neuron numbers are maintained in aging mouse distal colon. Neurogastroenterol Motil [Internet]. 2013 [cited 2026 Oct 22];25(7). Available from: https://pubmed.ncbi.nlm.nih.gov/23517051/

60. Tanaka H, Yamashita T, Asada M, Mizutani S, Yoshikawa H, Tohyama M. Cytoplasmic p21(Cip1/WAF1) regulates neurite remodeling by inhibiting Rho-kinase activity. J Cell Biol [Internet]. 2002 Jul 22 [cited 2026 Oct 22];158(2):321–9. Available from: https://pubmed.ncbi.nlm.nih.gov/12119358/

