## Supplementary figures for "Aging enteric neurons develop chromatin remodeling, DNA damage and a senescence-like phenotype"

### SUPPLEMENTARY DATA

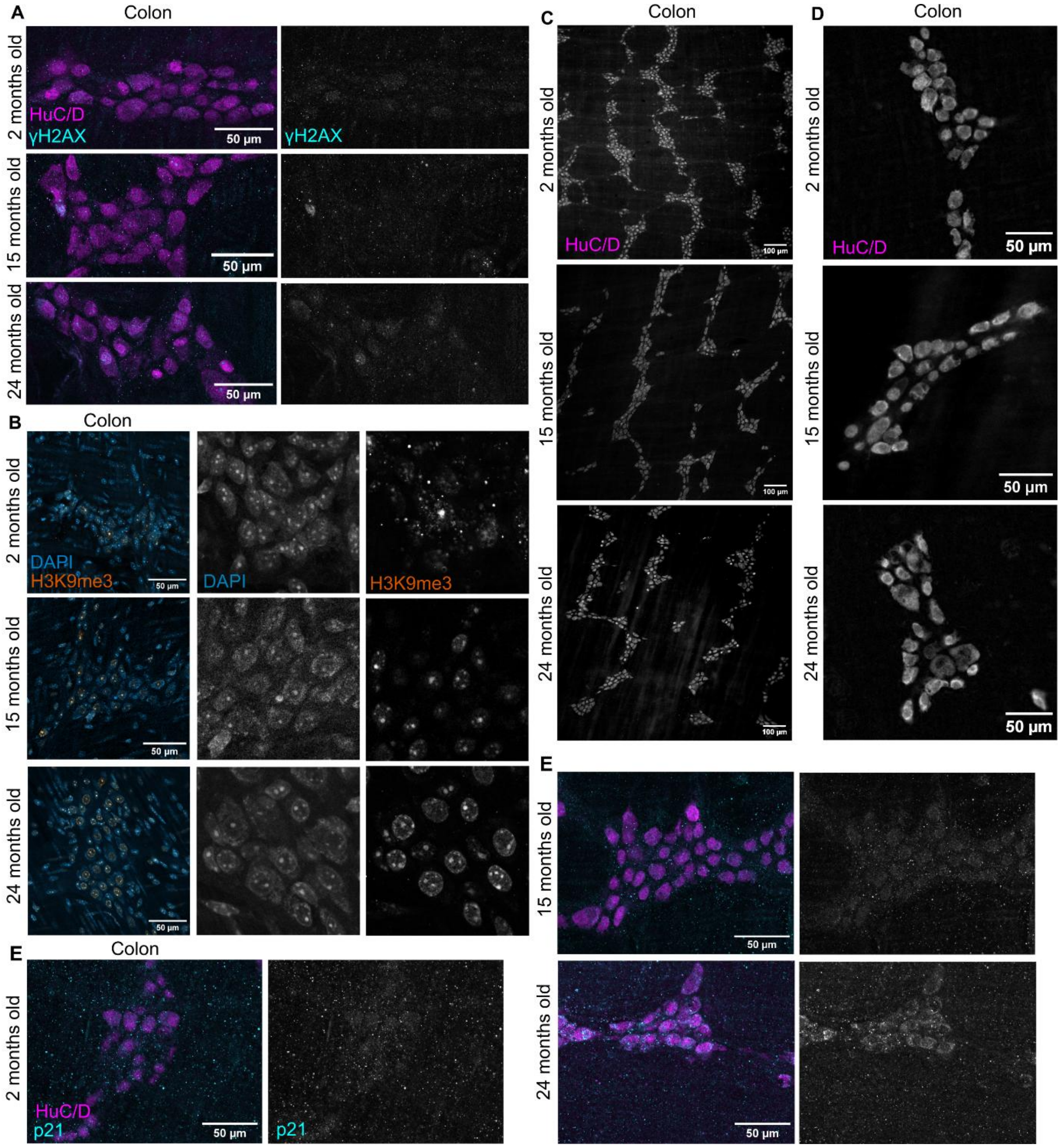


**Supplementary figure 1. Extended imaging data from figures 1 and 2.** (A) Representative γH2AX and HuC/D co-stainings in 2-, 15-, and 24-months old colonic myenteric plexus. (B) Representative images of in 2-, 15-, and 24-months old colonic myenteric ganglia immunolabeled for H3K9me3 and DAPI. (C) Representative HuC/D immunostainings of colonic LMMP preparations from 2-, 15-, and 24-months old mice for visualization of decreasing neuronal numbers with age. (D) Representative HuC/D immunostainings from 2-, 15-, and 24-months old colonic LMMPs used for neuronal area analysis. (E) Representative images of 2-, 15-, and 24-months old colonic myenteric ganglia immunolabeled for p21 and HuC/D.

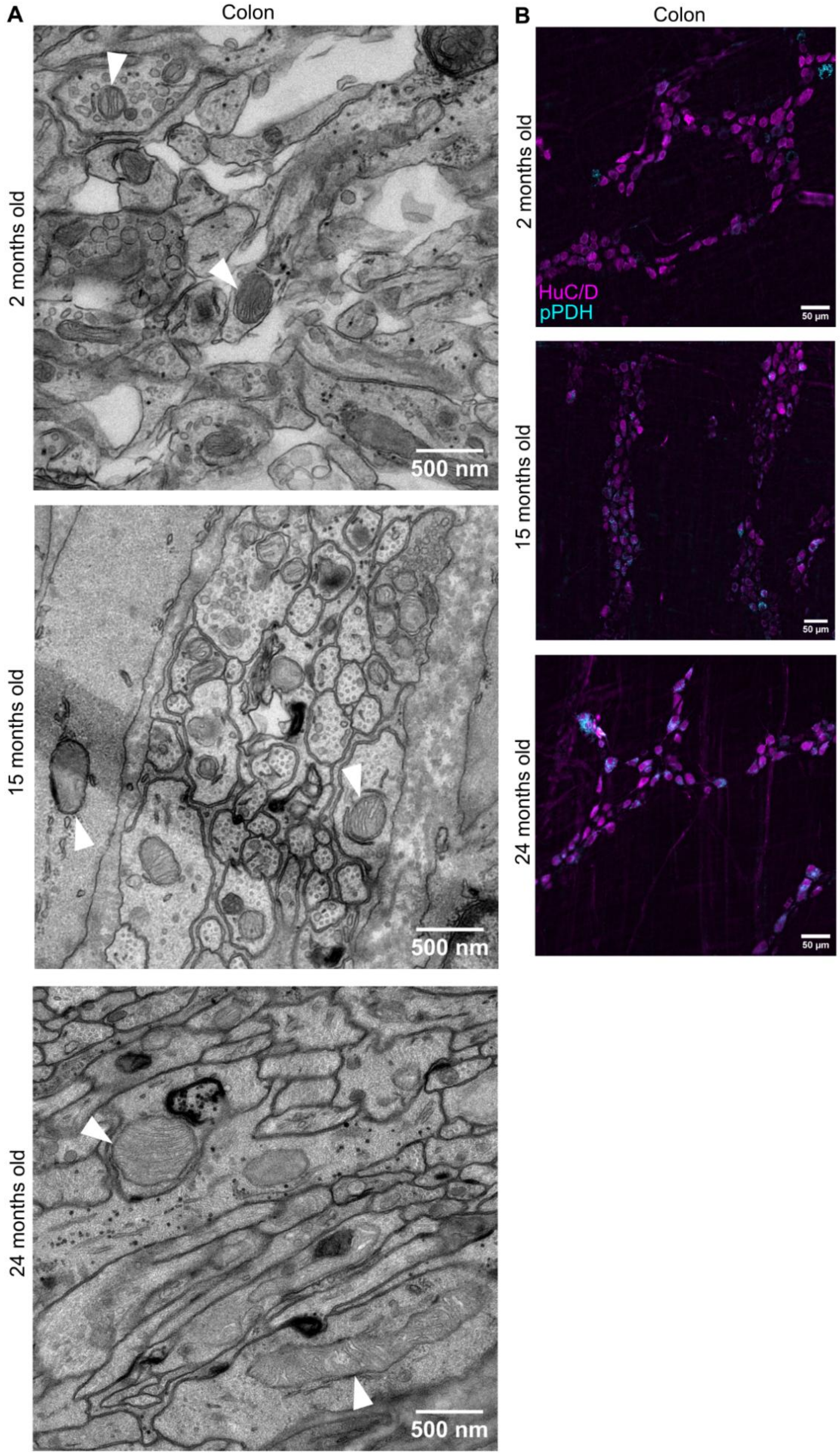

**Supplementary figure 2. Extended imaging data from figure 4.** (A) Representative transmission electron microscopy images of colonic LMMP cross-sections from 2-, 15-, and 24-months old mice. White arrowheads indicate mitochondria. (B) Representative images of 2-, 15-, and 24-months old colonic LMMPs immunolabeled for pPDH and HuC/D.
